# Mechanics and growth coordination define SOSEKI-based polarity fields

**DOI:** 10.1101/2025.11.26.690761

**Authors:** Marcel Piepers, Blanca Jazmin Reyes-Hernández, João R D Ramos, Neva Bölke, Laura Schütz, Changzheng Song, Anamarija Primc, Lotte Bald, Roeland MH Merks, Dolf Weijers, Alexis Maizel

**Affiliations:** Center for Organismal Studies (COS), University of Heidelberg, Im Neuenheimer Feld 230, 69120 Heidelberg, Germany; Mathematical Institute, University Leiden, P.O. Box 9512, 2300 RA, Leiden, The Netherlands; Institute of Biology, Leiden University, P.O. Box 9505, 2300 RA, Leiden, The Netherlands; Laboratory of Biochemistry, Wageningen University, Wageningen 6708WE, The Netherlands

## Abstract

The formation of organs requires the coordinated growth of cells and tissues relative to the main body axes. Plant growth is typically anisotropic and mechanically coupled through contiguous cell walls, yet how these physical patterns link to cell and tissue polarity remains unclear. Here, we use SOSEKI (SOK) proteins—previously thought to report global polarity fields—as markers to dissect how polarity arises during lateral root (LR) organogenesis. Live imaging revealed that SOK polarisation does not follow a uniform global field but instead responds to local differences in growth and mechanical state between neighbouring tissues. SOKs accumulate at interfaces separating domains of distinct growth behaviour and dissipate under compressive stress. After perturbing tissue interfaces or the mechanical continuity of the tissue, the coherence of the polarity was disrupted, indicating that stable axis establishment requires mechanical coupling across the cell wall network. Our results suggest that SOK polarisation is mechanoresponsive, linking tissue mechanics to polarity and axis establishment.

## Main Text

Multicellular development requires the coordinated growth of cells, tissues, and organs relative to the major body axes. In plants, organ growth is characterised by the preservation of contiguous cell wall contacts, is typically anisotropic, and produces specific shapes. While animals rely on canonical polarity proteins, these have not been identified yet in plants, which employ alternative strategies, including auxin transport, cell wall–mediated mechanical continuity, and geometry-based cues. The local orientations of anisotropic expansion, known as the principal directions of growth (PDGs), are inherently linked to organ shape and mechanics. PDGs provide a geometric framework for how growth patterns can inform division orientation and tissue architecture (Hejnowicz & Romberger, 1984). Yet, the molecular mechanisms that connect PDGs to division orientation remain largely unknown.

SOSEKI (SOK) proteins represent a deeply conserved family of polar proteins present throughout land plants. Their polarisation depends on an N-terminal DIX domain that mediates concentration-dependent polymerisation, enabling the formation of discrete polar domains at cell edges. This polymerisation is not merely structural: it provides the high local concentration required to recruit effector proteins such as *ANGUSTIFOLIA*, thereby linking polarity landmarks to cellular functions. Remarkably, DIX-dependent polymerisation is shared with animal polarity proteins such as Dishevelled (Schwarz-Romond *et al*, 2007) and SAR-group “Musketeer” proteins (Kostareli *et al*, 2025), underscoring an ancient and conserved biochemical principle of polarity establishment across kingdoms (van Dop *et al*, 2020). In *Arabidopsis thaliana*, SOKs are expressed in the embryo and proliferating regions of the primary and lateral roots (LRs), and have been proposed to interpret global polarity cues and influence cell division orientation locally (Yoshida *et al*, 2019). However, the nature of the cues that determine SOK localisation is unclear.

The post-embryonic formation of lateral organs, such as LRs, requires the respecification of growth and cell division along axes orthogonal to those of the main organ. The growth of these lateral organs is also characterised by unsteady growth fields, in which growth anisotropy varies in direction or between adjacent regions, potentially creating mechanical conflicts (Szymanowska-Pułka et al., 2012). It is unknown when and how the polarity fields are established upon the formation of the new organ axes. In *Arabidopsis*, LRs originate from xylem pole pericycle (XPP) cells that maintain cell proliferation in the differentiation zone and act as founder cells (Dubrovsky *et al*, 2001). LR development follows a stereotyped series of stages (Malamy & Benfey, 1997). Upon initiation (LRI), activated founder cells radially swell and divide asymmetrically to generate a packet of small central cells flanked by larger ones (Fig. 1a, Stage I; (Vermeer *et al*, 2014; Vilches Barro *et al*, 2019; Torres-Martínez *et al*, 2020)). These cells then divide periclinally, forming the two-layered Stage II primordium (Malamy & Benfey, 1997; von Wangenheim *et al*, 2016; Schütz *et al*, 2021) (Fig. 1a). Subsequent stages involve continued division and growth, progressively shaping the primordium into an organisation comparable to the primary root. During this process, anisotropic expansion predominates in the central domain (provascular domain), while the surrounding ground tissue expands less, resulting in continuous deformation of the primordium (Malamy & Benfey, 1997; von Wangenheim *et al*, 2016). In the primary root apex, whose geometry remains steady, SOK proteins localise to polar cell edges, faithfully reporting the shoot–root and radial axes that align with the principal directions of growth in a stable growth field (Nakielski, 2008). By contrast, in the emerging LR, SOKs polarise to edges oriented along the new shoot–root and radial axes, which are orthogonal to those of the parent root (Yoshida *et al*, 2019) (Fig. 1a). How SOK polarity is established and reoriented during the formation of new organ axes, and what cues determine this respecification, remains unknown. Here, we show that local mechanical and geometric cues, rather than a global field, define coherent SOK-based polarity fields in developing LRs.

**Figure 1.**
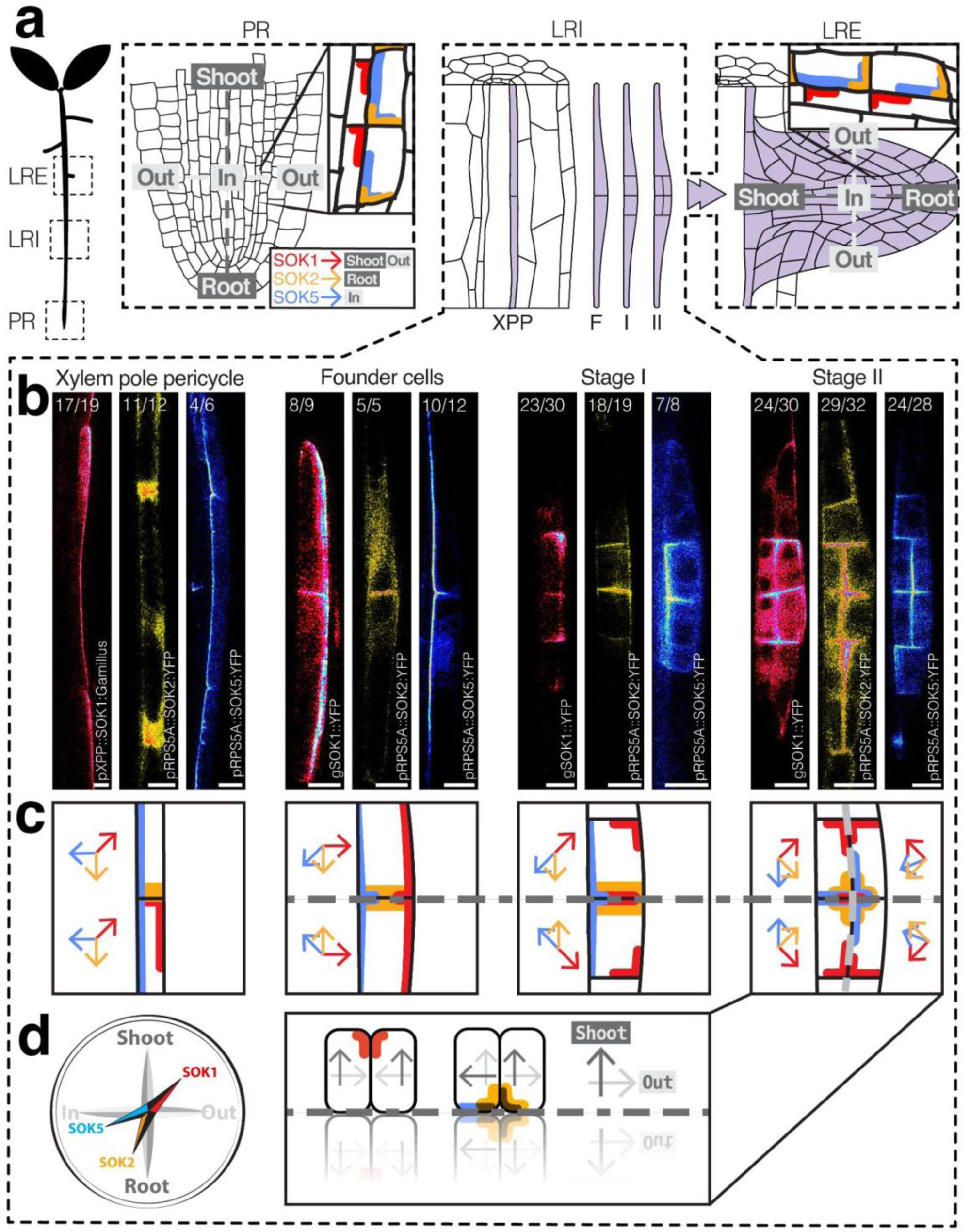
Two symmetry-breaking events pattern early lateral root primordia. **a**, Schematic of *Arabidopsis* root zones showing the primary root (PR), lateral root initiation (LRI), and lateral root emergence (LRE). Insets illustrate SOSEKI (SOK1 in red, SOK2 in orange, SOK5 in blue) localisation in the PR and emerged LR, reporting the shoot–root and in–out axes. **b**, Representative images of fluorescently tagged SOK1, SOK2, and SOK5 reporters in XPP cells, founder cells, Stage I, and Stage II primordia. Numbers indicate the frequency of observed patterns across independent samples. Scale bars, 10 µm. See also Movie S1. **c**, Consensus schematics of SOK polarisation at each stage. Arrows denote the orientation of SOK polarisation in each cell. The grey dashed lines indicate axes of symmetry. **d**, At Stage II, the shoot-root and in-out axes inferred from SOK polarisation are incoherent between the inner and outer cells.

To investigate the emergence of the SOK polarity fields during LR formation, we mapped the polarisation of SOK1, SOK2, and SOK5 from founder cells activation to LR emergence. Expressed from their native promoters, these three SOKs could all be detected during LR development, albeit with distinct expression strengths and domains (Yoshida *et al*, 2019). To ensure reliable tracking of polarity across all cells, we complemented these with RPS5A-driven reporters (Weijers *et al*, 2001) for SOK1, SOK2, and SOK5, as well as an XPP-specific (Vilches Barro *et al*, 2019) SOK1 reporter to assess pre-activation polarisation. Importantly, these misexpression constructs did not alter cell division orientation in the root meristem or LR development, and polarisation remained unperturbed (Fig. S1a, b). These validated reporters enabled us to track the polarisation of SOK1, SOK2, and SOK5 from activation to Stage II (Fig. 1a–c, Fig. S2). In XPP cells, all three proteins polarised as in the primary root meristem, integrating the parent root polarity field (Fig. 1b XPP; schematic in Fig. 1c). Upon founder cell activation and radial swelling, however, their polarisation diverged: SOK1 shifted to the edges away from the founder-founder interface and to the periclinal face toward the endodermis, SOK2 concentrated at the first cell wall formed after the founder cell division, and SOK5 decorated edges flanking this interface and the vasculature-facing periclinal edge (Fig. 1b, founder cells). After the first asymmetric division (Stage I), SOK1’s divergent polarisation became even more pronounced (Fig. 1b, Stage I, schematic in Fig. 1c). These changes mark the first symmetry-breaking event, where diverging polarity fields replace the parental SOK pattern of XPP centred around the central cell wall. A second symmetry-breaking occurred during the transition to Stage II, when periclinal divisions created a two-layered primordium. Here, SOK1 polarized toward the new periclinal interface between inner and outer cells, SOK2 formed a cross-like pattern marking both the first cell wall formed after the founder cell division and periclinal interfaces, and SOK5 likewise accumulated at the periclinal interface (Fig. 1b, Stage II; schematic in Fig. 1c). Flanking cells that had not yet divided retained their earlier polarisation, highlighting the domain-specific reorganisation of polarity (Fig. S2).

Because SOK polarisation in mature roots and embryos defines a polarity “code” for the shoot–root and in–out axes (Yoshida *et al*, 2019), we next asked whether the early symmetry-breaking events already establish these axes (Fig. 1d). In mature roots, endogenous SOK1 marks the shoot-out edge, while SOK2 and SOK5 localise to the root-in edge. At Stage II, however, their axial patterns were inconsistent: SOK1 converged toward the new periclinal interface, whereas SOK2 and SOK5 diverged away from it (Fig. 1d). Thus, although both shoot-root and radial information can be detected, the combined SOK patterns cannot be reconciled into a coherent set of shoot–root and radial axes. These observations demonstrate that, early on, SOK proteins do not obey a uniform global polarity field, suggesting that the establishment of the LR axis requires a later maturation phase.

To understand how incoherent polarity patterns become progressively organised, we tracked the polarisation of SOK1, SOK2, and SOK5 during stages III–VI of LR development (Movie S1). Strikingly, despite the *pRPS5A* promoter driving uniform expression throughout LR development (Fig. S3), the pRPS5A::SOK1/2/5:YFP signal was not evenly distributed. While polarisation was stable in individual cells, it differed between tissue domains (Fig. 2a). Fluorescence intensity was consistently enriched near the interface separating provascular and ground tissue precursors, corresponding to the prospective pericycle–endodermis (PE) interface (Fig. 2a, Movie S2). To quantify this, we processed light-sheet and confocal datasets spanning stage I to emergence using a custom pipeline that identified the positions of 1,388 SOK signal maxima across 462 images, yielding a density map of SOK signal maxima. This analysis confirmed that all three SOK reporters preferentially accumulated near the PE interface throughout LR organogenesis (Fig. 2b). Interestingly, each SOK displayed a slightly different localisation bias. pRPS5A::SOK1:YFP enriched toward ground tissue interfaces, consistent with its known attraction to the endodermis–cortex junction in the root apical meristem (Yoshida *et al*, 2019). pRPS5A::SOK2:YFP accumulation aligned more precisely with the PE interface itself, while pRPS5A::SOK5:YFP was enriched in provascular cells directly adjacent to the PE interface (Movie S1). Together, these patterns suggest that the PE interface constitutes a consistent attractor for SOK polarisation, with each family member responding to slightly differently within this region. Interestingly, the PE interface, which corresponds to the periclinal interface specified upon the Stage I to II transition (Fig. S4, Movie S2), separates the anisotropically growing provascular core from the more isotropic ground tissue. Quantification of cell anisotropy across stages confirmed that growth anisotropy progressively increased in the central provascular domain. In contrast, growth in the ground tissue precursors remained isotropic (Fig. 2c). Importantly, SOK accumulation did not directly correlate with anisotropy at the per-cell level (Fig. S5). Instead, SOKs accumulate near the tissue boundary where domains of differential growth meet (Fig. 2b).

**Figure 2.**
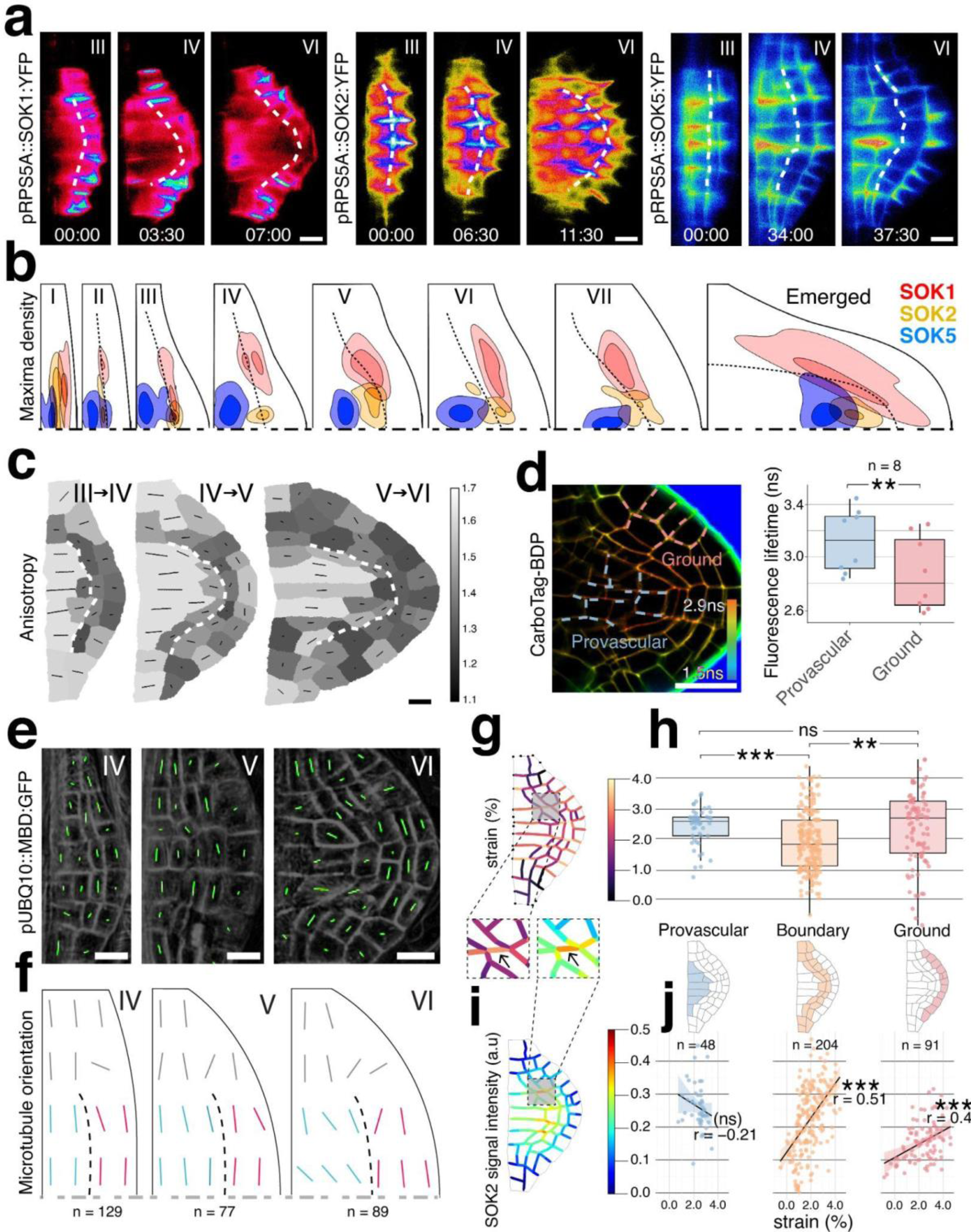
SOK polarisation highlights domains of differential tissue growth. **a**, Time-lapse light-sheet imaging of SOK1, SOK2, and SOK5 reporters from LR primordium stages III to VI. SOK polarisation is stable in individual cells and is consistently enriched near the pericycle–endodermis interface (dashed line). Scale bars, 10 µm. Time in hh:mm format. **b**, Consensus density maps highlighting the regions with the highest SOK1 (red), SOK2 (orange), and SOK5 (blue) expression across developmental stages. Developmental stages and number of cells analyzed: I (*n* = 126), II (*n* = 216), III (*n* = 294), IV (*n* = 234), V (*n* = 122), VI (*n* = 69), VII (*n* = 114), emerged (*n* = 213). Range of LR height at each stage: I: 8-15μm, II: 15-23μm, III: 23-30μm, IV: 30-38μm, V: 38-45μm, VI: 45-50μm, VII: 50-60μm, emerged: 60-130μm. Only half of the LR primordium is represented. The black dashed line is the pericycle-endodermis interface. **c**, Quantification of cell growth anisotropy between successive stages (III–IV, IV–V, V–VI). Higher anisotropy is detected in the central domain. Scale bar, 10 µm. **d**, Left, confocal FLIM imaging of CarboTag-BDP in stages III–V of LR primordia. Color scale denotes fluorescence lifetime (ns). Right, quantification of CarboTag-BDP fluorescence lifetimes in epidermal, ground tissue, pericycle-endodermis junction, and provascular domains. Provascular and pericycle show significantly higher values (**p<0.01). **e**, Confocal imaging of cortical microtubule organisation (pUBQ10::MBD:GFP, grey) at stages IV–VI. Microtubule orientation and anisotropy in each cell are indicated by green overlays. Scale bars, 10 µm. **f**, Schematics of average microtubule orientation. Central tissues show alignment along anisotropic growth directions, while peripheral tissues remain isotropic. The black dashed line is the pericycle-endodermis interface. **g**, Predicted strain pattern obtained after pressurising the cellular template of a Stage V primordium. **h**, Boxplots showing the distribution of strain across the provascular, boundary, and ground tissue domains. **i**, SOK2 cortical signal intensity mapped on the same pressurised template as in g. **j**, Correlation between strain magnitude and SOK2-YFP signal intensity for each tissue domain.

To test whether the domains of differential growth correspond to differences in mechanical state, we next analysed cell wall properties and cytoskeletal organisation across the developing primordium. Cell wall fluorescence lifetime imaging with the carbohydrate-binding dye CarboTag-BDP (Besten *et al*, 2025), <u>2025),</u> reporting cell wall network structure, did not reveal evident spatial heterogeneity in wall environments of primordia Stages I to V (Fig. S6). However, in later stages, lifetime quantification revealed significantly longer lifetimes in central domains than in peripheral ones, suggesting differences in cell wall composition or stiffness (Fig. 2d). This result echoes similar conclusions from the primary root (Alonso Baez *et al*, 2025). We then examined cortical microtubule organisation as a proxy for mapping the orientation of mechanical stresses (Verger *et al*, 2018; Vilches Barro, 2018) in developing LR primordia (Fig. 2e). Quantification of the mean anisotropy-weighted microtubule orientation at cell-level regions revealed a supracellular pattern of microtubule orientation (Fig. 2f). In the central domain of stages IV and V, the MT orientation is perpendicular to the PDG, whereas in stage VI, the MT orientation in the provascular domain changed and tended to align with the PDG of the provascular domain, reflecting anisotropic growth in the provascular domain. In contrast, peripheral cells displayed more isotropic microtubule arrays, consistent with their reduced and directionally unconstrained expansion. Together, these data demonstrate that the LR primordium is mechanically heterogeneous: the provascular domain is characterised by denser walls and anisotropic stresses, whereas the surrounding ground tissue has less dense walls and isotropic stresses. Thus, the provascular-ground tissue interface emerges as a boundary between domains of differential growth and distinct mechanical states. This suggests that SOK polarisation may be conditioned by local mechanical heterogeneity rather than a uniform global polarity field.

To further explore the relationship between tissue mechanics and SOK polarisation, we developed a mechanical model of the developing LR primordium (Fig. S7a). The model was based on segmented experimental data from *pRPS5A::SOK2* reporter lines at stages IV–VI and was used to calculate local strain distributions across the provascular, ground tissue, and their intervening boundary zone. Importantly, the simulations did not impose any predefined differences in cell-edge properties between regions, allowing the mechanical response to emerge solely from tissue geometry that resulted from growth patterns. The simulated strain distribution revealed lower strain values in the boundary region between the provascular and ground tissue domains (Fig. 2g-i, Fig. S7b, c), demonstrating that tissue geometry alone is sufficient to generate mechanical heterogeneity within the primordium and highlighting this boundary as a mechanically distinct region. We then examined the correlation between SOK2 signal intensity at individual cell edges and the corresponding local strain magnitude (Fig. 2g, h, j, Fig. S7d). We observed a moderate, albeit significant, correlation between the SOK signal and local mechanical strain at cell edges in the boundary and, to a lesser extent, in the ground tissue regions, suggesting a link between polarity and the mechanical landscape of the developing primordium.

Building on the observation that SOKs map to domains of differential growth and mechanical state, we next asked how polarity responds when these tissues are subjected to changes in mechanics. LR emergence provides a natural context for this as primordia successively push through overlying tissue layers (Vilches-Barro & Maizel, 2015). The endodermis acts as a mechanical barrier, generating compressive stress at stage II (Stöckle *et al*, 2022). While the cortex and epidermis constrain the shape of the LR primordium (Lucas *et al*, 2013), it remains to be established whether this occurs through compressive stress. To address this, we ablated cortex cells overlying a stage IV primordium and observed an immediate increase in primordium length (7µm in 5 seconds, Fig. S8a). This rapid elongation indicates that cortical cells exert compressive stress. We next asked how SOK polarisation responds to these stresses. Time-lapse imaging of pRPS5A::SOK5:YFP from stage II to emergence revealed a striking correlation between SOK accumulation and growth (Fig. 3a; Movie S3). SOK5 signal intensity increased during periods of elongation but sharply decreased during phases of little or no growth, despite the *RPS5A* promoter being active throughout organogenesis (Fig. S8b). The decline in signal coincided with developmental windows during which the primordium encounters maximum compressive stress beneath the endodermis (stage II) and the cortex (stage IV). A similar pattern was observed for pRPS5A::SOK2:YFP, suggesting that dissipation under stress is a general feature of SOK proteins (Fig. S8c). To confirm this interpretation, we relieved compressive stress by ablating overlying cortex cells and imaged SOK5 signal dynamics. In ablated samples, the SOK5 signal remained consistently high throughout organogenesis, with intensities approximately 3.5-fold greater than in controls (Fig. 3b, Movie S3). To more precisely address the relationship between changes in tissue mechanics induced by cell ablation and SOK polarisation, we extended our mechanical model to simulate strain distribution in lateral root primordia following removal of the overlying cortex and epidermis (Fig. 3c). Using the same mechanical parameters as for intact primordia, the ablation model revealed a significant increase in strain values, and an increased heterogeneity (larger variance) of strain values in periclinal walls compared to non-ablated controls (Fig. 3d). These simulations demonstrate that releasing external compression through ablation shifts mechanical load toward periclinal interfaces. Interestingly, this shift is consistent with experimentally observed enhancement of SOK5 polarisation along these surfaces (Fig. 3e). Together, these results suggest that the mechanoresponsive behaviour of SOKs reflects their ability to sense and align with the redistribution of tensile forces within the primordium in response to mechanical perturbation. If correct, a reduction of the tensile forces due to increased compressive forces should lead to a reduction in SOK signal accumulation. To experimentally test this prediction, we perturbed microtubules in the endodermis using the inducible driver–reporter system *pCASP1»PHS1ΔP* to trigger endodermal microtubule depolymerisation, leading to a lack of endodermis thinning and an increase in compressive stresses (Stöckle *et al*, 2022). Dexamethasone (DEX) treatment triggered endodermal microtubule depolymerisation, verified by a marked loss of

**Figure 3.**
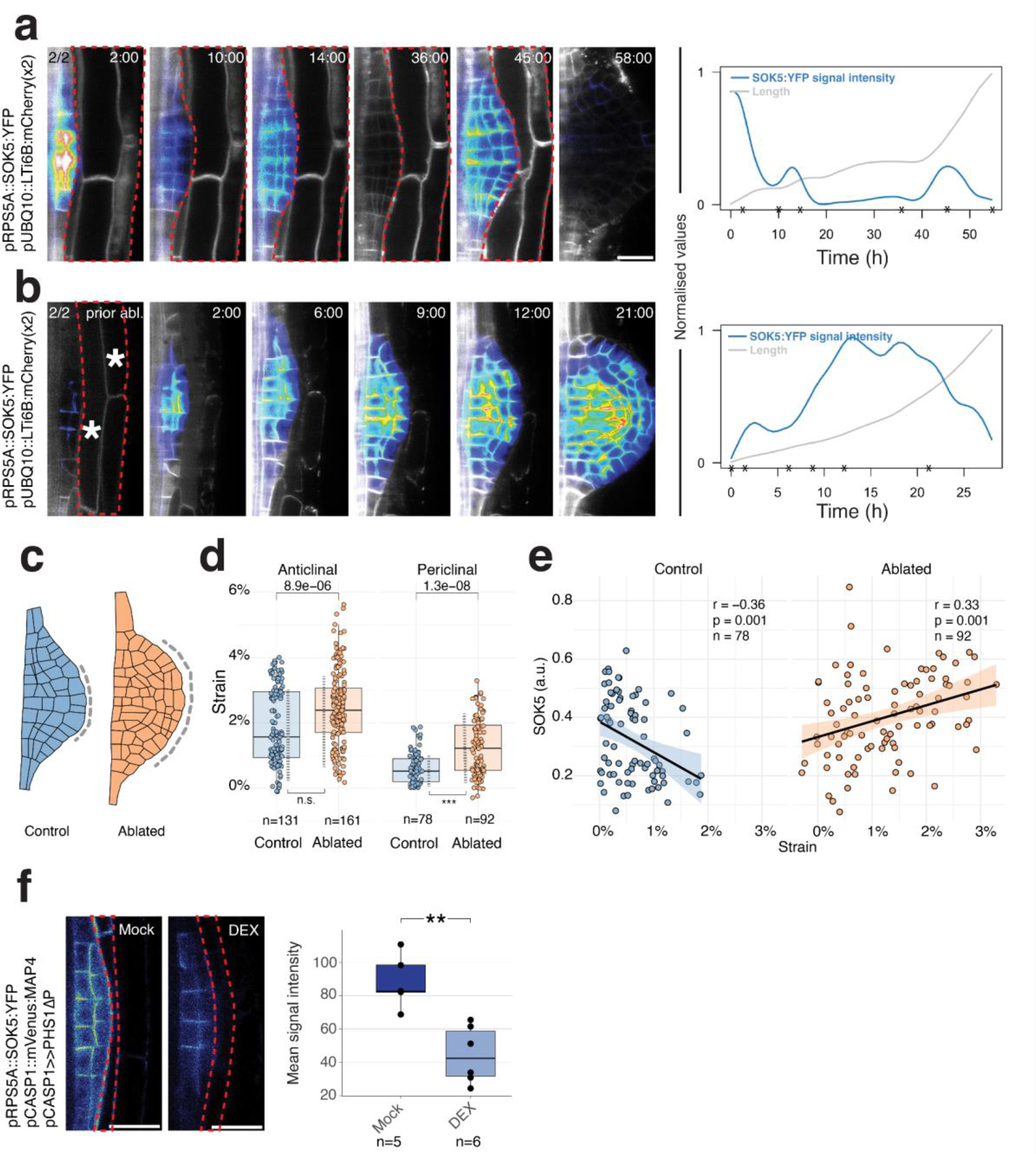
SOSEKI accumulation is mechanoresponsive during lateral root emergence. **a, b** Time-lapse light-sheet imaging of pRPS5A::SOK5:YFP (blue) with plasma membrane marker (grey) from stage II to emergence in wild type (a) and after targeted ablation of overlying cortex cells (asterisks) to relieve compressive stress. Right: quantification of primordium length (grey) and SOK5 signal intensity (blue) over time. **c**, schematic of the meshes used for turgor-induced strain simulation in control and ablated conditions. The dashed lines indicate the range of external vertices set free to move. In the ablated case, the fraction of free-to-move vertices is increased compared to the control condition to account for the absence of the overlying tissue. **d**, boxplot showing the distribution of strain in anticlinal and periclinal walls after pressurising the cellular template in control and ablated conditions. P-values for the comparisons of the mean are indicated above. The vertical dashed lines indicate the distributions’ variances and the symbols below indicate the result of a Levene’s test for equality of variance (***, p<0.001). Anticlinal walls are perpendicular to the primordium surface, periclinal ones parallel. **e**, Correlation between strain magnitude and SOK5-YFP signal intensity in periclinal walls in control and ablated conditions. **f**, Confocal imaging of pRPS5A::SOK5:YFP upon dexamethasone (DEX)-inducible depolymerisation of endodermal microtubules using pCASP1»PHS1ΔP. Scale bars, 20 µm.

MAP4:mVenus signal (Fig. 3f, Fig. S8d). In this background, pRPS5A::SOK5:YFP intensity was significantly reduced compared to mock controls, indicating that compressive stress generated by defective endodermal responses dissipates the SOK signal. Together, these results establish that SOK proteins are polarity markers whose accumulation is mechanoresponsive, linking the physical constraints of emergence to the dynamic regulation of polarity fields in the LR primordium.

To test whether the mechanoresponsive behaviour of SOKs extends beyond the root, we examined their localisation in the shoot apical meristem (SAM), an organ where tissue dynamics generate heterogeneous stress patterns (Alonso-Serra *et al*, 2024; Shi *et al*, 2018; Hamant *et al*, 2008; Uyttewaal *et al*, 2012). Despite uniform activity of the *pRPS5A* promoter (Fig. S9a), pRPS5A::SOK1/2/5:YFP signals were specifically absent from creases near the peripheral zone and at sites of incipient primordia (Fig. S9b). These regions are characterised by minimal growth and elevated mechanical stress arising from organ outgrowth and boundary formation. Thus, as in LR primordia, SOKs dissipate from mechanically stressed zones in the SAM, suggesting that their stress response is a general feature of plant organogenesis rather than an organ-specific phenomenon.

Our previous results showed that SOK polarity reflects mechanical heterogeneity and is mechanoresponsive. We next asked how cell and tissue interfaces, as well as mechanical continuity, contribute to the establishment and maintenance of SOK polarity patterns. We first live-imaged gSOK1::YFP in response to the targeted ablation of the overlying endodermis at Stage II. Removal of the endodermis resulted in a loss of edge polarity in the adjacent outer central cells (Fig. 4a), a phenotype reminiscent of natural endodermal thinning during stage II, which also coincides with reduced SOK1 polarity (Fig. 4b). These results indicate that the endodermis is required not only as a structural contact but may also serve as a conduit for transmitting mechanical cues that stabilise SOK1 polarity at adjacent cell edges. We then asked whether the cell wall network within the primordium is also playing a role in SOK1 polarity. For this, we performed single-cell ablations in stage I and II primordia (Fig. 4c). Of 24 ablation events, half led to an immediate abortion of the primordium development. In contrast, the others continued for some time but eventually arrested. In this group, the SOK1 signal was frequently lost, although transient polarity reorientations were observed near the ablation site (Fig. 4c, Fig. S10a). These results indicate that targeted alteration of the cell wall network is sufficient to reconfigure polarity fields and alter organ-level behaviour. Because SOK polarisation maps to the provascular–ground tissue boundaries, where domains of differential growth and distinct mechanical states meet, we next asked whether altering root radial patterning would affect SOK polarisation. To this end, we examined SOK1 polarity in the *scarecrow* (*scr)* mutant, which is defective in proper ground tissue patterning and fails to establish the outer layer during the stage II–III transition (Goh *et al*, 2016). In wild type, SOK1 polarity gradually shifts from the PE interface toward the ground tissue interface during stages II–IV. In *scr* mutants, however, polarity was disrupted throughout the primordium, including provascular cells normally unaffected by *SCR* function (Fig. 4d). This indicates that proper patterning and differentiation of the ground tissue are required for long-range polarity coherence. We can speculate that in the *scr* mutant, changes in the mechanical properties and/or the topology of the tissue cell wall network alter the transmission of mechanical cues, stabilising SOK polarity across the primordium.

**Figure 4.**
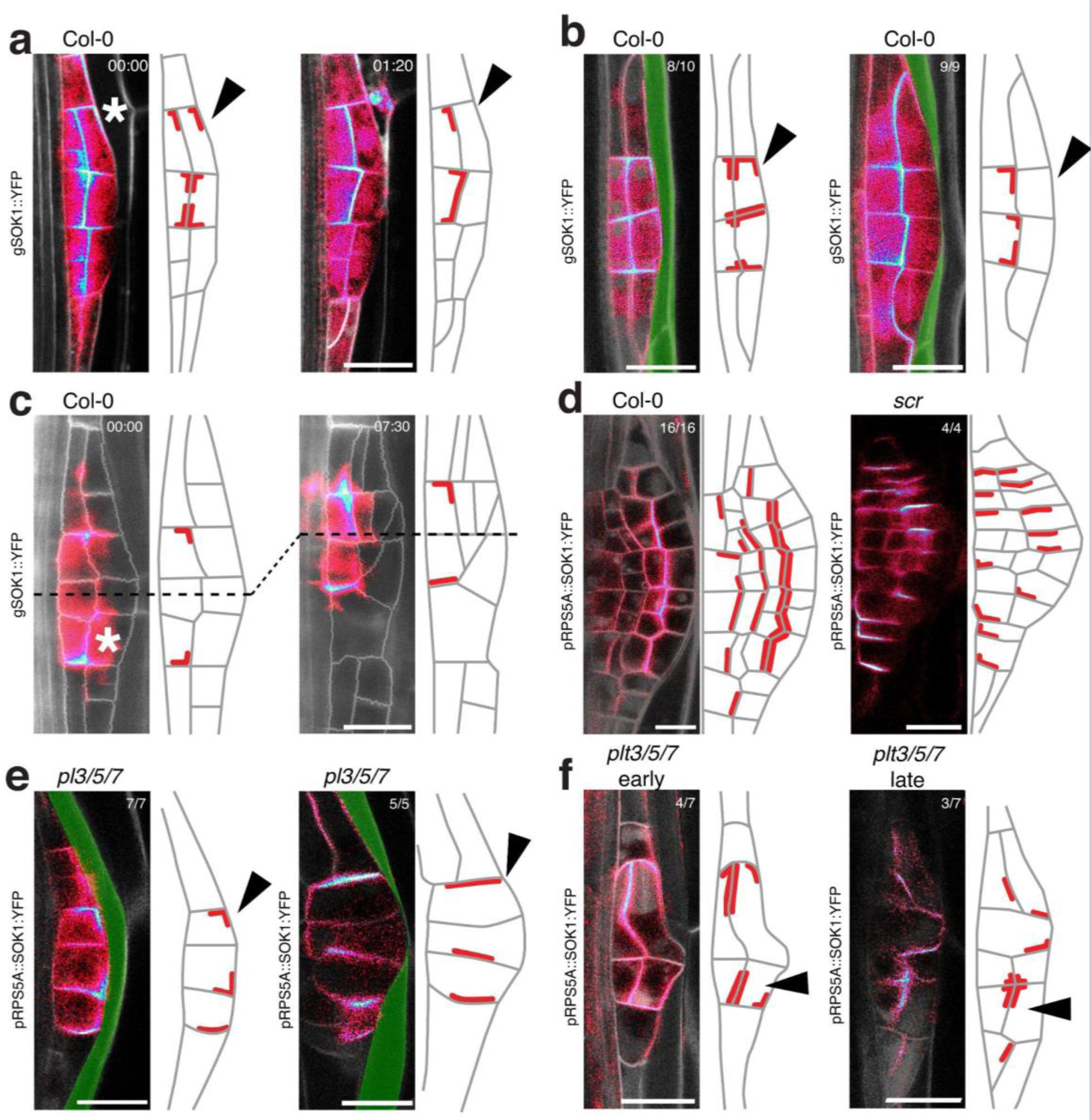
SOK polarity depends on tissue interfaces and mechanical continuity. **a-f**, Confocal or light-sheet (c) sections through LR primordia expressing the indicated SOK reporter. In each case, the images and matching schematic depict the consequence of the following perturbations on SOK polarisation: (a) ablation of endodermis, (b) crossing of endodermis, (c) ablation of a cell in the LR primordium, (d) *scr* mutant, (e) crossing of the endodermis or (f) formation of a periclinal interface in *plt3/5/7* mutant. Asterisks mark ablated cells. Black arrowheads highlight changes in polarisation. Scale bars: 20 µm.

We next focused on the role of the PE interface, which depends on the PLETHORA3/5/7 (PLT3/5/7) transcription factors (Du & Scheres, 2017). In *plt3/5/7* triple mutants, the PE interface is rarely formed, yet SOK1 maintained its divergent edge polarity in stage I primordia unless the overlying endodermis was thinned, reiterating that the endodermis-primordium interface affects SOK1 polarity (Fig. 4e). Remarkably, in rare *plt3/5/7* primordia that underwent a periclinal division in some cells, SOK1, SOK2, and SOK5 repolarised to this new interface, resembling the wild type pattern (Fig. 4f early, Fig. S10b). This wild-type-like pattern was absent in older primordia (Fig. 4f early, Fig. S10b), which were characterised by more isotropically oriented cortical microtubules than in wild type and by deformed cells growing isotropically (Fig. S10d). Together, these data suggest that, while the transient establishment of a correct cellular layout can account for proper SOK polarisation independent of specific cell identity, changes in growth behaviour lead to SOK repolarisation. To investigate when mispolarisation occurred, we recorded pRPS5A::SOK2:YFP in *plt3/5/7* and found that mispolarisation coincided with growth arrest, confirming the mechanosresponsiveness of SOK accumulation (Fig. S10d). Together, these data show that SOK polarity depends on the structural presence of cell wall networks and their mechanical continuity. Disruption of interfaces, whether by ablation or genetic perturbation, redistributes mechanical stresses and alters polarity at the organ level. In contrast, perturbations in growth or geometry that preserve cell–cell contacts typically retain wild-type-like polarity. Thus, the mechanical continuity of the cell wall network emerges as a key determinant of SOK polarity during organogenesis.

Together, our results reveal that SOK proteins provide a molecular link between growth fields and axis formation in plants. Rather than reflecting a uniform global field, SOK polarity emerges from the physical architecture of tissues—interfaces, boundaries, and the continuity of the cell wall network. These findings establish SOKs as sensitive reporters of mechanical organisation in living tissues and highlight how the coupling between mechanics and polarity underpins the self-organisation of plant organs. Future work should aim at clarifying SOSEKI proteins are direct mechanosensors or act downstream of mechanical signal transduction as well as elucidating the developmental functions of SOKs. This will be key to understanding how plants integrate mechanical information to coordinate growth, patterning, and morphogenesis.

## Methods

### Plant material and growth conditions

All experiments were performed in *Arabidopsis thaliana* (Col-0 background). An overview of plant lines used is provided in Supplementary Table S1. SOSEKI marker lines (*gSOK1–5::YFP*) were described previously (Yoshida *et al*, 2019) and were crossed with plasma membrane marker lines (pUBQ10::LTI6B:mCherry; (Elsayad *et al*, 2016)). SOK reporters were introgressed into the *plt3/5/7* and *scr* mutants backgrounds (Hofhuis *et al*, 2013; Goh *et al*, 2016), by transformation or crossing. Seeds were surface-sterilised (70% ethanol, 0.05% SDS), stratified for 24 h at 4 °C in darkness, and sown on half-strength Murashige and Skoog (½MS) medium (pH 5.7) supplemented with 1% phytoagar. Plants were grown under long-day conditions (16 h light/8 h dark, 22 °C). For some experiments, plates were supplemented with 1% sucrose. Soil-grown plants were cultivated under the same photoperiod and temperature.

### Plant transformation and selection

Stable transformation was performed using the floral dip method (Clough & Bent, 1998) with *Agrobacterium tumefaciens* strain GV3101. Transformants were selected either by antibiotic resistance (BASTA, hygromycin, kanamycin) or by fluorescence-accumulating seed technology (FAST; (Shimada *et al*, 2010)).

### Inducible expression

For endodermis-specific microtubule depolymerisation, we used the *pCASP1>>PHS1ΔP* system (Stöckle *et al*, 2022). Seedlings were transferred at 7 d after germination (DAG) to ½MS medium containing 15 μM dexamethasone (DEX) and imaged after ∼24 h.

### Sample preparation and staining

Seedlings were fixed in 4% paraformaldehyde in PBS for 2 h at room temperature or overnight at 4 °C, then washed and cleared using ClearSee (Kurihara *et al*, 2015). Cell walls were counterstained with 1 mg/mL calcofluor white. For live imaging, seedlings were mounted in ½ MS medium on agar slices, in chambered coverglass systems, or in glass/FEP capillaries for light-sheet microscopy.Confocal and light-sheet microscopy Confocal imaging was performed on Leica SP5II and SP8 systems equipped with water-immersion objectives and white-light or multi-photon lasers. Excitation/emission settings are provided in Supplementary methods. For live imaging, z-stacks were acquired every 30 min with step sizes of 0.5–1 μm. Light-sheet imaging was performed using a Luxendo MuViSPIM (Bruker) with dual-sided illumination and 40× or 60× water-immersion objectives. Time-lapse recordings spanned up to three days, with z-stacks collected at 30-minute intervals. Additional details are provided in the Supplemental Materials.

### Laser ablation

Targeted ablations were performed either with two-photon excitation on a Leica SP8 system (960 nm, 20–30% power) or with an integrated UV ablation laser on the MuViSPIM. Exposure times were adjusted until visible damage was achieved. Additional details are provided in the Supplemental Materials.

### Image processing and quantification

An extensive description is provided in the Supplemental Materials. Images were processed in Fiji. Light-sheet data were corrected and cropped using BigDataProcessor2 (Tischer *et al*, 2019). Drift correction of confocal timelapses was done with the Fiji Registration plugin. For weak signals, maximum intensity projections of 2–10 slices were generated. SOK signal maxima were quantified using a custom Python pipeline on 462 images (1388 maxima) and plotted relative to developmental stage. CarboTag-BDP fluorescence lifetimes were fitted with a two-component exponential decay model in LAS X software (Leica) and analysed in R. Microtubule orientation was quantified from 2D cortical images using FibrilJ (Boudaoud *et al*, 2014), with anisotropy-weighted mean orientations plotted per cell group. Growth anisotropy and principal growth directions were determined with MorphoGraphX (Strauss *et al*, 2022). Additional details are provided in the Supplemental Materials.

### Molecular cloning

All constructs (listed in Table S1) were generated by GreenGate cloning (Lampropoulos *et al*, 2013; Piepers *et al*, 2023).

## Data availability

All raw images, processed data, and biological resources are available from the corresponding author upon reasonable request. Custom scripts for image analysis are available from https://doi.org/10.5281/zenodo.17598920.

## Acknowledgements

We thank Joseph Dubrovsky for his critical reading of the manuscript. Research in the Maizel lab is supported by the German Science Foundation (DFG) through the Research Unit FOR2581 (project P06). J.R. and R.M.H.M. were supported by the Dutch Research Council (NWO) through the Gravitation program GreenTE (project no. 024.006.001).

## Ethics declarations

The authors declare no competing interests.

## Contributions

A.M. D.W. and M.P. conceived the study and analyzed the results. M.P. performed wet bench experiments under the supervision of A.M. and, with help from or material generated or provided by B.J.R.H, N.B., L.S. C.S., A.P., L.B. nd D.W.. J.R.D.R. performed the computational modelling with input from A.M. and R.M.H.M, who provided feedback on interpretation. M.P. and A.M. wrote the manuscript with input from all authors.

