## Supplemental Material for "Mechanics and growth coordination define SOSEKI-based polarity fields"

Piepers et al.

### Content


- **Figure S1**. Heterozygous ectopic expression of SOK1, SOK2 and SOK5 does not alter LR development and polarisation.
- **Figure S2**. Similar SOK polarisation between independent samples during early LR organogenesis.
- **Figure S3**. Homogeneous RPS5A promoter activity during lateral root organogenesis.
- **Figure S4**. The pericycle-endodermis interface from stage II to the emerged lateral root.
- **Figure S5**. Low correlation between cortical SOK1/2/5 signal intensity and anisotropic cell growth at the single cell level.
- **Figure S6**. CarboTag-BDP fluorescent lifetimes reveal no prominent differences in cell wall structure in developmental stages from XPP to stage V.
- **Figure S7**. Correlations between strain and SOK2 in tissue domains across developmental stages.
- **Figure S8**. SOK2 accumulation tracks growth dynamics.
- **Figure S9**. SOK signal is absent in creases of the shoot apical meristem.
- **Figure S10**. SOK polarity post ablation and in the plt3/5/7 mutant.

- **Movie S1**. Lightsheet microscopy time-lapse recording of SOK1, SOK2 and SOK5 expressed from the pRPS5A promoter
- **Movie S2**. Pericycle-endodermis interface tracking during lateral root development
- **Movie S3**. pRPS5A::SOK5:YFP signal during wildtype-like emergence and unobstructed emergence.

- **Supplemental Methods**
- **Supplemental Table**. Resources used and generated
- **References**

### Supplemental Figures

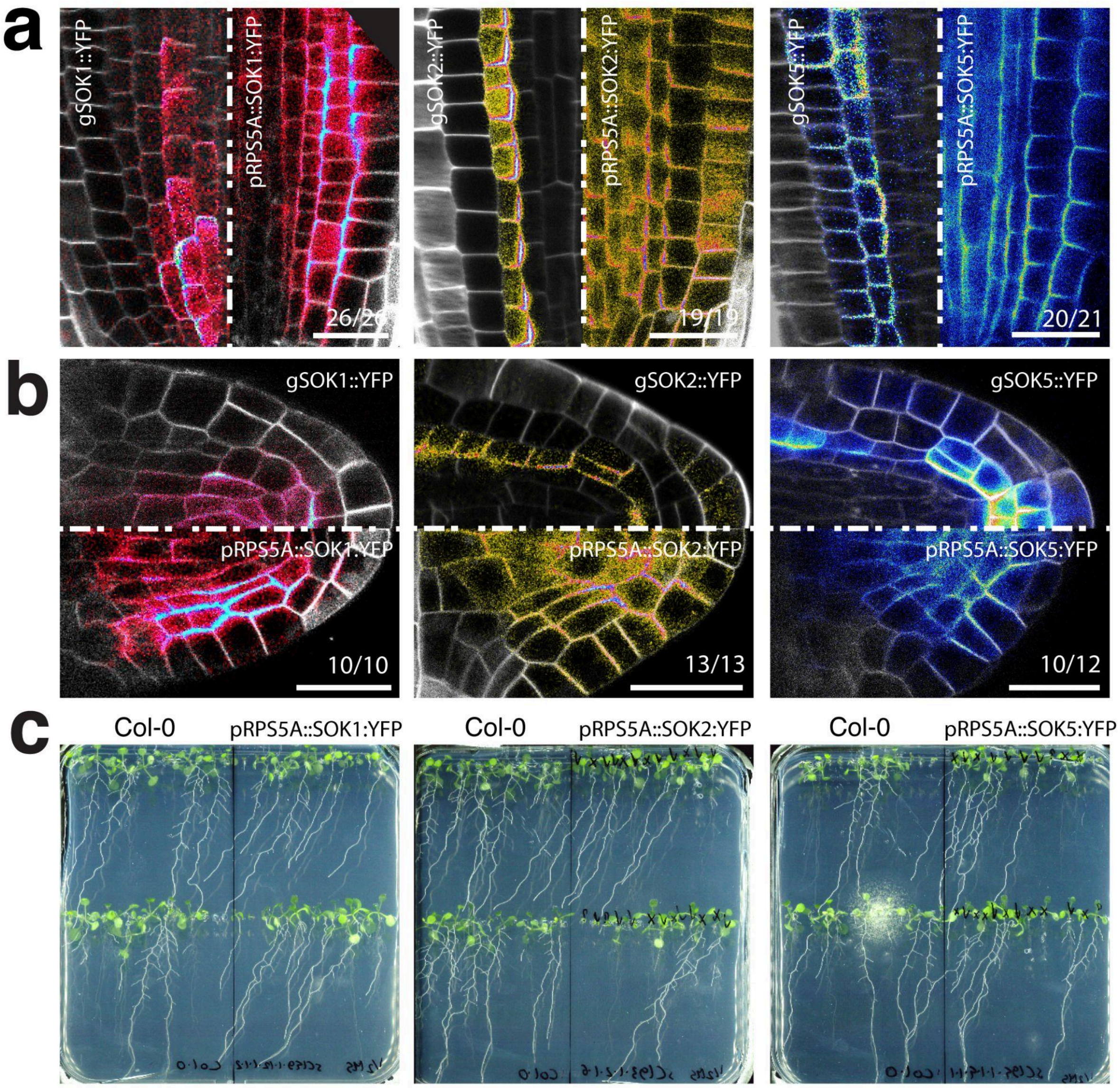


**Figure S1 | Heterozygous ectopic expression of SOK1, SOK2 and SOK5 does not alter LR development and polarisation.**

**a,** Representative images of the root apical meristem in heterozygous *pRPS5A::SOK1/2/5* lines show no noticeable disruption to cell layout.

**b,** Emerged LRs in the same lines display normal tissue organisation. Numbers in the lower right of each panel indicate the penetrance of the observed pattern. Cell contours are marked by pUBQ10::LTI6B:mCherry(x2). Scale bar: 20µm.

**c.**,Root system of 7-day-old seedlings grown on ½MS plates for wildtype (Col-0) and indicated SOK misexpression lines.

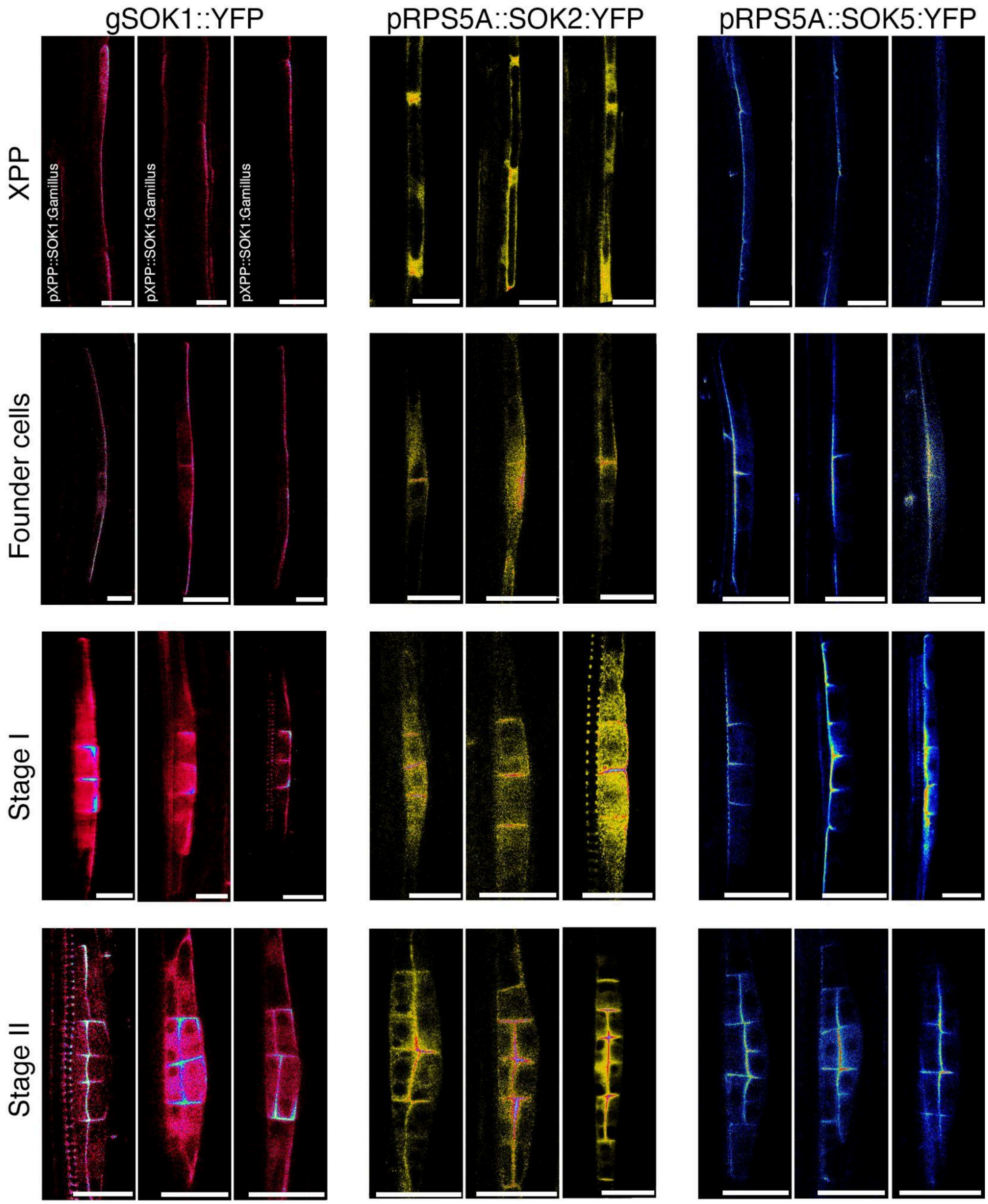


**Figure S2 | Similar SOK polarisation between independent samples during early LR organogenesis**

Microscopy images of SOK markers during early lateral root organogenesis show similar polarisation between independent samples for each developmental stage. Images were obtained using confocal or multiphoton microscopy, on fixed or live samples. Importantly, the polarity pattern is independent of imaging method or fixation treatment. The SOK polarity patterns are visualised with the indicated markers. Scale bar: 25μm.

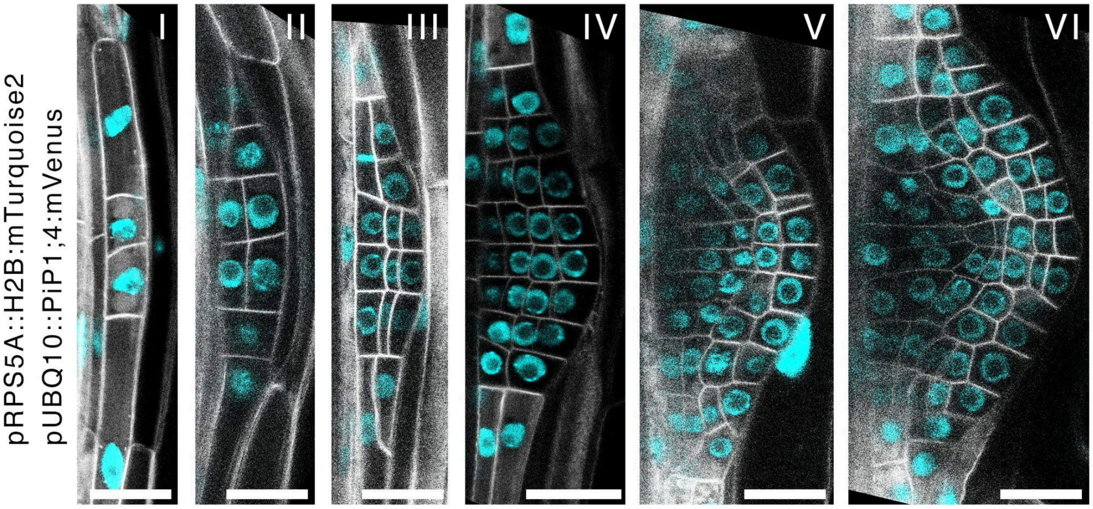


**Figure S3 | Homogeneous *RPS5A* promoter activity during lateral root organogenesis**
pRPS5A::H2B:mTurquoise2 signal (cyan) is visible in all cells throughout LRP development. Scale bar: 20μm.

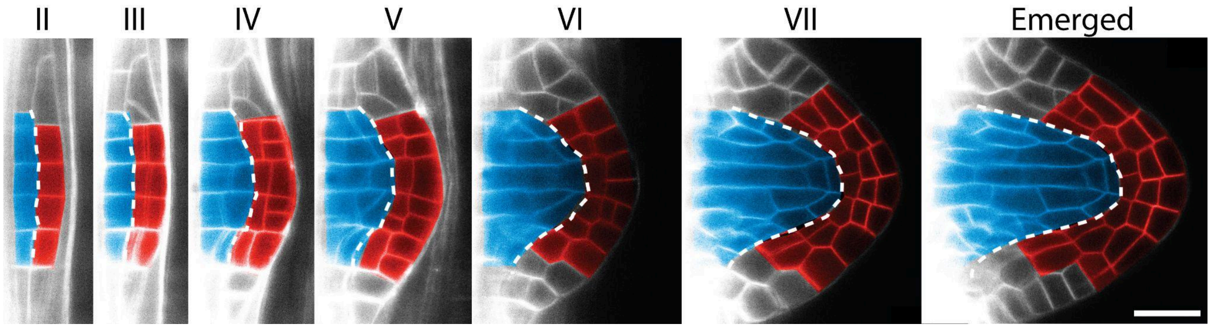


**Figure S4 | The pericycle-endodermis interface from stage II  to the emerged lateral root.** Lightsheet microscopy time-lapse of pUBQ10::LTI6B:mCherry(x2) plasma membrane marker. The provascular (blue) and ground tissue precursors (red) are colored digitally. The future pericycle-endodermis interface (dashed white line) lies at the interface of the provascular and ground tissue precursors. Scale bar 20µm. See also Movie S2.

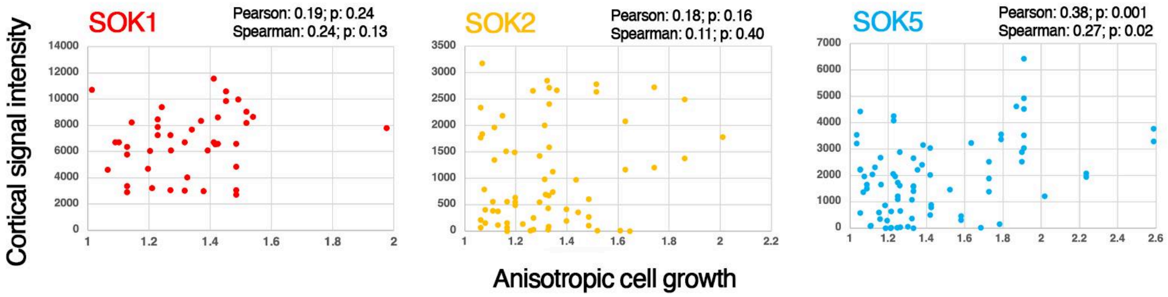


**Figure S5 | Weak correlation between cortical SOK1/2/5 signal intensity and anisotropic cell growth at the single cell level.**

Scatter plots of cortical signal for the indicated SOK protein as a function of the anisotropy in cell growth of a given cell.

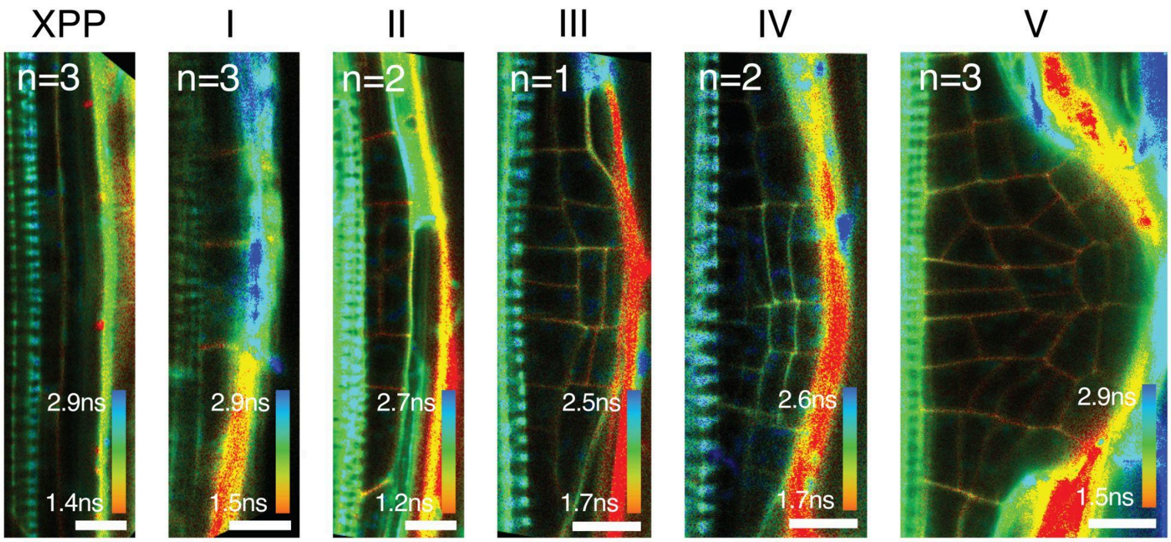


**Figure S6 | CarboTag-BDP fluorescent lifetimes reveal no prominent differences in cell wall structure in developmental stages from XPP to stage V.**

FLIM-based imaging of LRP at the indicated developmental stage stained with CarboTag-BDP. Scale bar: 20 µm

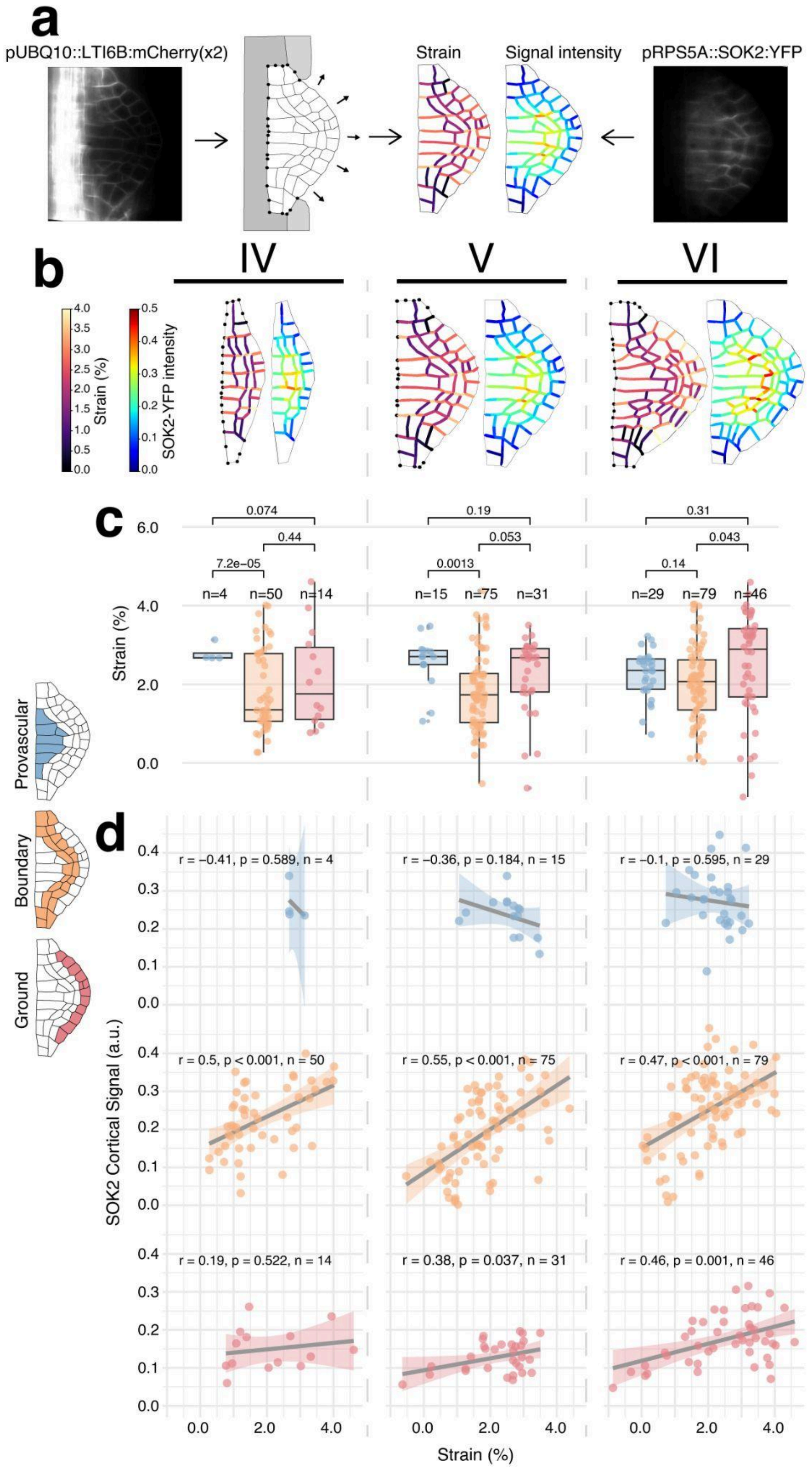


**Figure S7 | Correlations between strain and SOK2 in tissue domains across developmental stages.**

**a**, Workflow for extracting primordium geometry, predicting strain, and quantifying SOK2-YFP signal intensity. The tissue mesh was manually segmented from the plasma membrane marker (pUBQ10::LTI6B:mCherry(x2)). The model was pressurised under a defined turgor pressure, with internal and adjacent layer vertices (dark grey) fixed and selected surface vertices (light grey) constrained to mimic mechanical resistance from neighbouring cells. The resulting strain field of the simulated primordium was computed. The experimental SOK2-YFP signal was mapped onto the same mesh, and fluorescence intensity was quantified along cell edges using Gaussian convolution.

**b**, Predicted strain (left) and corresponding SOK2-YFP signal (right) patterns in LRP at stages IV–VI.

**c**, Boxplots showing the distribution of strain across the provascular, boundary, and ground tissue domains at the indicated stages.

**d**, Correlation between strain magnitude and SOK2-YFP signal intensity for each tissue domain for the indicated stage.

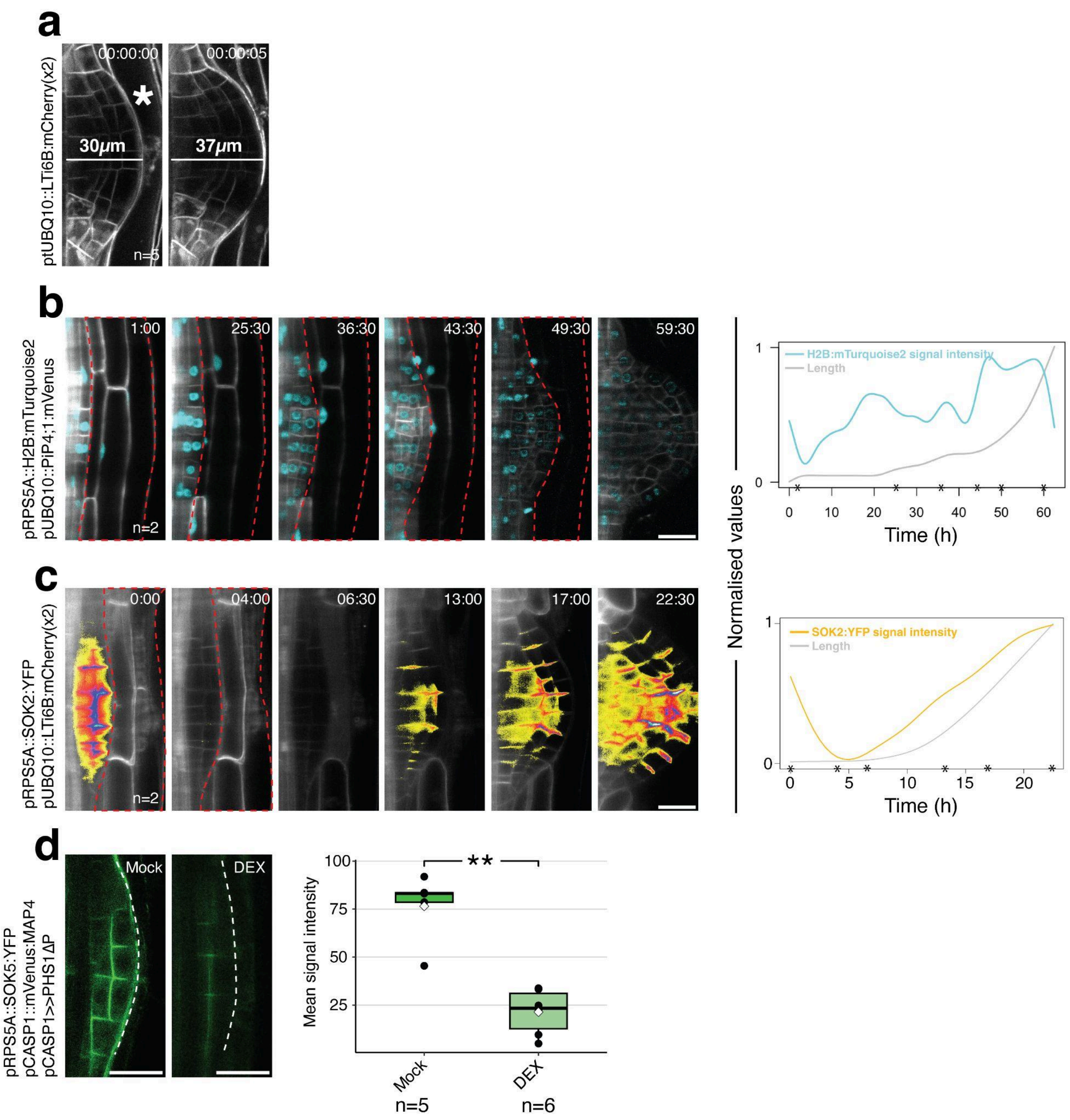


**Figure S8 | SOK2 accumulation tracks growth dynamics**

**a**, Laser ablation of cortical cells (asterisk) in stage IV primordia expressing the plasma membrane marker pUBQ10::LTI6B:mCherry(x2) results in an immediate increase in lateral root width, consistent with rapid release of mechanical compression. Time in hh:mm:ss format.

**b**, Time-lapse imaging of pRPS5A::H2B-mTurquoise2 and pUBQ10::PIP1;4:mVenus during lateral root growth shows that *RPS5A* promoter activity (mean normalised signal intensity ratio, mTurquoise2/mVenus) does not correlate with primordium elongation (right).

**c**, In contrast, pRPS5A::SOK2:YFP signal intensity (mean normalised YFP/mScarlet ratio) positively correlates with primordium elongation over time (right), indicating that SOK2 accumulation reflects growth dynamics rather than transcriptional activity. Time in hh:mm format. **d**, Endodermis-specific depolymerisation of microtubules using *pCASP1>>PHS1ΔP* leads to reduced cortical pCASP1::MAP4:mVenus signal intensity (dashed line), confirming microtubule depolymerisation following dexamethasone (DEX) treatment. Scale bars, 20 μm.

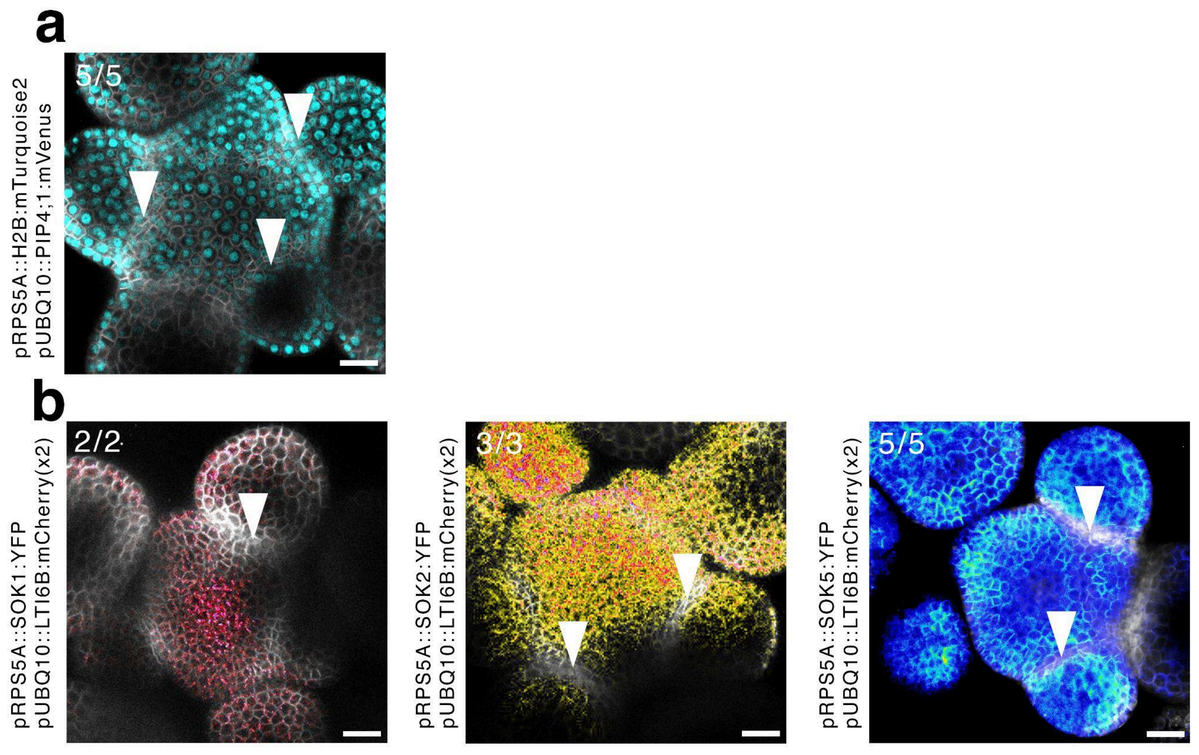


**Figure S9 | SOK signal is absent in creases of the shoot apical meristem**

**a**, **b**, Confocal microscopy image of shoot apical meristems expressing the indicated reporters from the *RPS5A* promoter. Numbers in the upper right corner indicate the penetrance of the depicted pattern. Arrowheads indicate the crease region between the SAM and the lateral organ primordia. Scale bar 20μm.

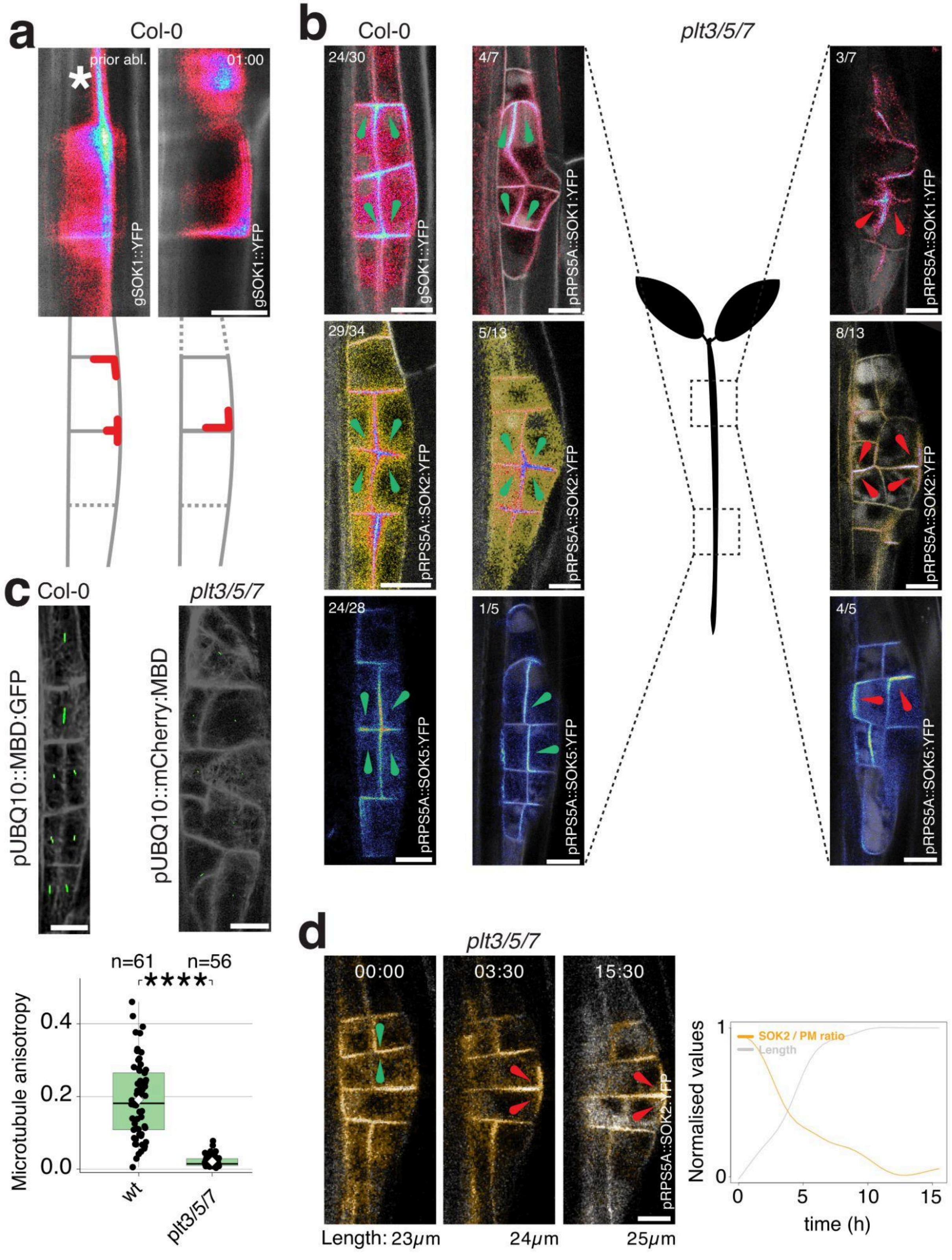


**Figure S10 | SOK polarity post ablation and in the *plt3/5/7* mutant.**

**a**, Light sheet microscopy imaging of SOK1 re-polarisation in response to ablation of a peripheral cell in stage I. The asterisks (*) indicates the ablated cell. The schematic below the microscopy images illustrates the repolarisation.
**b**, Confocal images of the indicated SOK reporter in wild type (Col-0) or *plt3/5/7* mutant. The SOK polarity pattern can be wildtype-like or aberrant in the *plt3/5/7* mutant. The mispolarized primordia were located in the upper (older) section of the primary root. In contrast, primordia with a WT-like polarity pattern were located in the lower section of the root.
**c**, Confocal time-lapse imaging of SOK2 in *plt3/5/7*. Quantification of SOK2 signal at the cell edge (normalised to the plasma membrane signal) shows that repolarisation of SOK2 coincides with a lack of growth. Green arrow heads indicate wt-like pattern of SOK polarisation, while the red ones highlight non-standard polarisation.
**d**, Images of MT orientation (green line) as determined by FibrilJ, length of the green line indicates anisotropy. Microtubule anisotropy is significantly lower in *plt/3/5/7* mutant Stage II-like primordia compared to wildtype primordia (****$p<0.0001$). Scale bars 10µm.

### Supplemental Movies

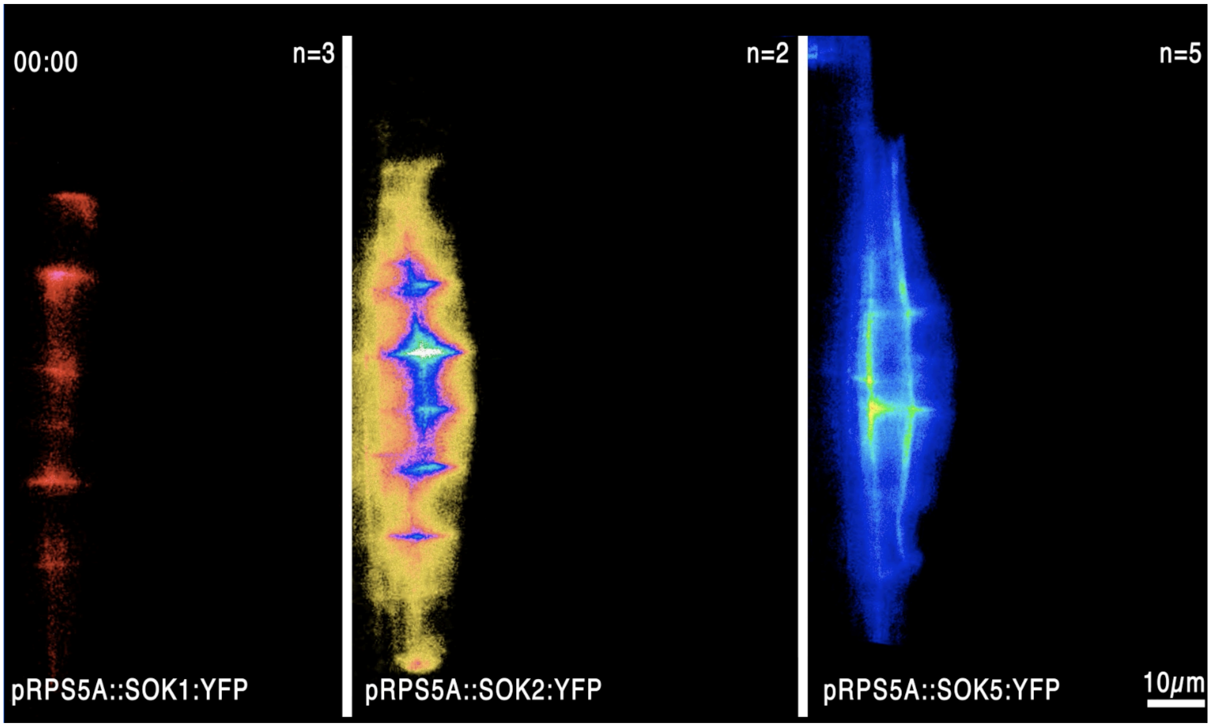


**Movie S1 | Lightsheet microscopy time-lapse recording of SOK1, SOK2 and SOK5 expressed from the pRPS5A promoter.**

Movie available: https://heibox.uni-heidelberg.de/f/faf2bf2ac9c24c5da310/

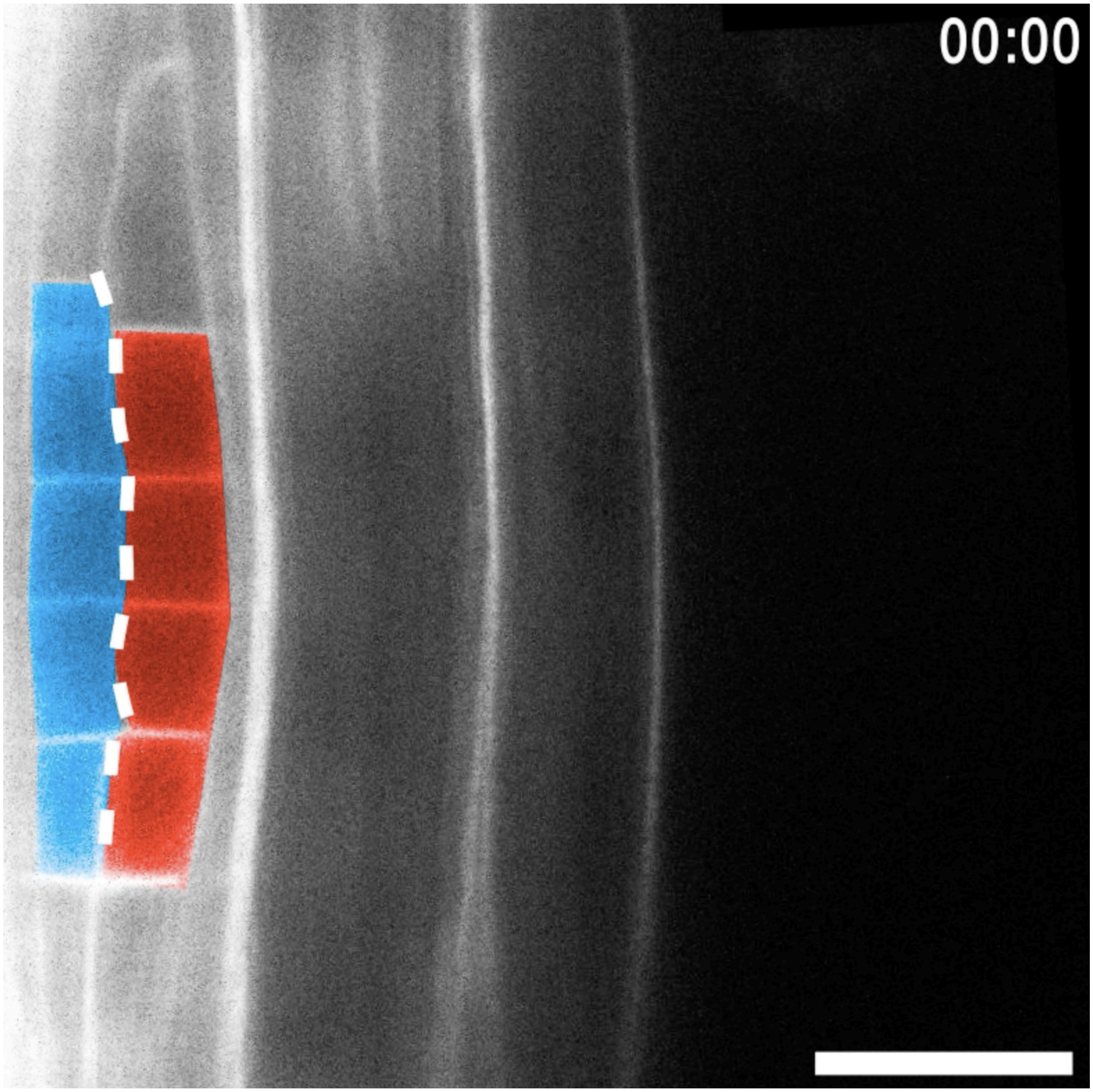


**Movie S2 | Pericycle-endodermis interface tracking during lateral root development**

Movie available: https://heibox.uni-heidelberg.de/f/01bf856ed34548e3a18d/

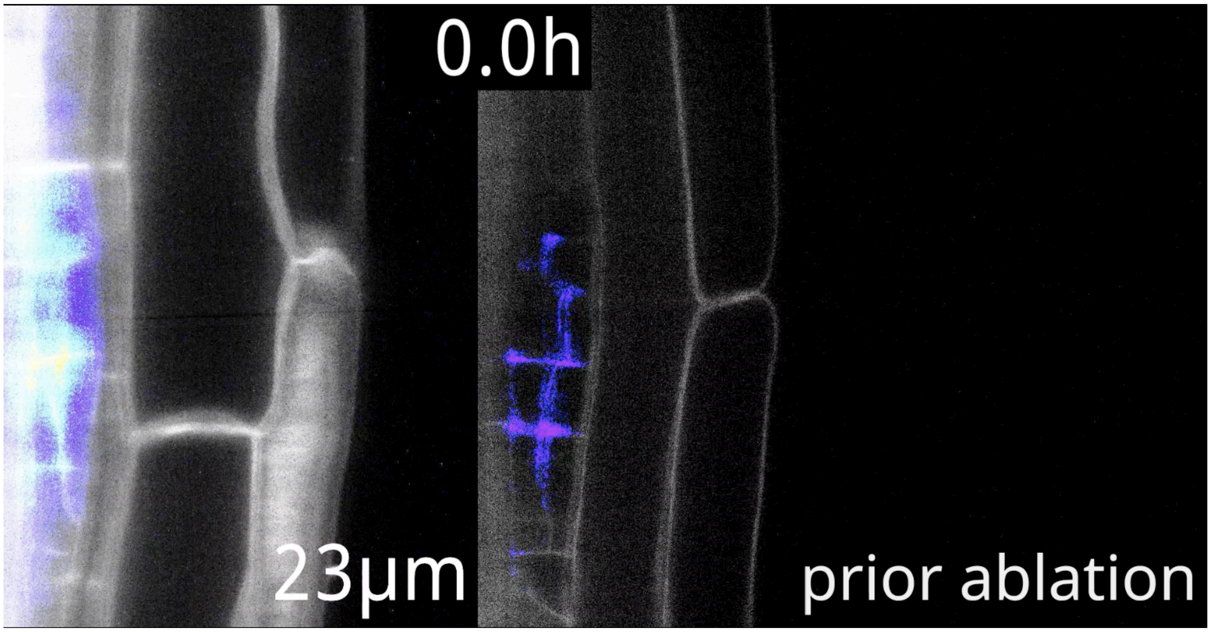


**Movie S3 | pRPS5A::SOK5:YFP signal during wildtype-like emergence and unobstructed emergence.**
https://heibox.uni-heidelberg.de/f/e845439be3a144febf99/

### Supplemental Methods

#### Sample Preparation and Imaging

#### Polarity observations in early LR organogenesis

For static analyses, confocal images and time-lapse snapshots were collected, marking SOK1, SOK2, and SOK5 polarity during pre-initiation (XPP), founder cell activation, stage I, and stage II primordia. Consensus polarity patterns were sketched as schematic cartoons. Polarity domain orientations (shoot, root, in, out) were interpreted relative to the root apical meristem polarity reference. Excitation/emission settings are provided below.

| Fluorophore | Excitation (nm) | Excitation two-photon (nm) | Emission (nm) |
|---|---|---|---|
| Calcufluor white | 405 | | 410-450 |
| mTurquiose2 | 458 | | 470-500 |
| GFP | 488 | 970 | 500-530 |
| Gamillus | 488 | | 500-530 |
| sYFP2 (YFP) | 514 | 970 | 520-550 |
| mVenus | 514 | | 520-550 |
| mScarlet-I | 561 | | 580-630 |
| mCherry | 561 | 1060 | 600-650 |

#### Light-sheet microscopy

Seedlings were mounted in glass capillaries (44.5 mm × 1 mm, 1.7 mm outer diameter) or fluorinated ethylene-propylene (FEP) tubes (44.5 mm × 1 mm, 1.6 mm outer diameter) filled with ½MS medium supplemented with Phytagel (8 mg/mL). Imaging was performed on a Luxendo MuViSPIM (Bruker) with dual-sided illumination. Detection was with a Nikon NIR Apo 40× NA=1.2 water-immersion objective, with an additional 1.5× lens to achieve 60× magnification for most timelapses. Fluorescent proteins were excited using diode lasers (458, 514, 561 nm). Emission was filtered through band-pass filter cubes (BP458–500 nm, 520–543 nm, 610–628 nm) and recorded simultaneously at opposing angles with sCMOS cameras. Z-stacks of 100 µm

(0.5 µm step size) were acquired every 30 min for up to 72 h. The dataset with the highest z-resolution for the region of interest was selected for further analysis. Timelapses were manually drift-corrected in Fiji using BigDataProcessor2 (Tischer *et al*, 2019).

**Laser ablation**

For ablation of the endodermis (Fig 4a), a two-photon excitation module of a Leica Stellaris SP8 setup was exploited to induce targeted cellular damage. The laser was set to 960nm and operated at 20–30% power, focused on a zoomed-in (40x) region encompassing a single endodermal cell. The exposure duration, approximately 1 second, corresponded to the frame acquisition time when using the following settings: 600 Hz scan speed, 512×512 px resolution, single frame per line, no frame averaging, and bidirectional scanning disabled.

For ablation of the overlying tissues or internal ablation (Fig 3b, 4c), a Luxendo MuViSPIM light sheet microscope (Bruker) equipped with an UV ablation laser was used (High Q, TDK-LAMBDA, Z36-12 REV:2.210 controller). The output power was approximately 1.8W. Ablation was performed via point illumination at the centre of the target cell. Exposure times were adjusted until visible damage was achieved.

### Data Analysis

**SOSEKI maxima density plots**

Timelapses (SOK1: n=3; SOK2: n=2; SOK5: n=5) and static confocal images (SOK1: n=21; SOK2: n=18; SOK5: n=8) were used to quantify LR polarity patterns. Preprocessing was done in Fiji: a 6 µm maximum projection at the primordium centre was rotated to orient LR growth to the right and cropped to consistent dimensions. A 25-pixel radius median filter was applied, converted to 8-bit, and thresholded to exclude background.

A custom script (Determine-SOK-maxima.py - available via https://doi.org/10.5281/zenodo.17598920) measured LR width (number of pixel columns with mean intensity > 0) and identified the three brightest SOK maxima iteratively, excluding previously assigned maxima. For static images, maxima were determined using the Find Maxima process in Fiji. Outputs were jointly plotted using Plot-SOK-Maxima.py (https://doi.org/10.5281/zenodo.17598920) . LR stages were assigned based on mean LR width: stage I (8–15 µm), II (15–23 µm), III (23–30 µm), IV (30–38 µm), V (38–45 µm), VI (45–50 µm), VII (50–60 µm), and emerged (60–130 µm). Final dataset sizes: I (n=126), II (n=216), III (n=294), IV (n=234), V (n=122), VI (n=69), VII (n=114), emerged (n=213).

**Growth anisotropy and principal growth directions**

Light-sheet timelapses from stage II–emergence (n=5) were analysed with MorphoGraphX (Barbier de Reuille *et al*, 2015). Plasma membrane channels were segmented with PlantSeg (Wolny *et al*, 2020) or manually in Photoshop. Cell meshes were constructed and used to calculate anisotropy tensors and principal directions of growth (PDGs). Domain-specific anisotropy was visualised as heatmaps overlaid on cell meshes.

**Supracellular microtubule orientation analysis**

Two-dimensional images of the cell cortex were acquired by positioning the image plane at the cell cortex using the two-photon module on a Leica Stellaris SP8 microscope. The microtubule fluorescence channel was processed in Fiji using the following workflow. Background subtraction using the rolling ball algorithm (radius = 100 pixels). Gaussian blur (σ = 1) to reduce noise. Contrast enhancement (saturation = 0.3, equalised) to improve visibility of microtubules. Unsharp masking (radius = 2 pixels, mask weight = 0.6) to enhance microtubule edges. Processed images were rotated to produce a rightward view of the LRP and cropped to match the measured LRP width.

Microtubule orientation and anisotropy were quantified for cells using manually drawn ROIs and the batch-processing version of FibrilJ (Boudaoud *et al*, 2014; Louveaux & Boudaoud, 2018). The resulting measurements were manually grouped according to developmental stage and the relative X and Y position of the corresponding cell within the LRP in Microsoft Excel. The relativel X and Y positions were corrected for difference in zoom level between images. X and Y coordinates in the bottom half of the primordia were mirrored to pool to the datapoints from both primordia halves.

The mean anisotropy-weighted microtubule orientation was calculated for measurement in Microsoft Excel using the following procedure. The measured orientation (in degrees) was converted to radians and doubled to account for the fact that orientations 180° apart correspond to the same microtubule orientation. For each measurement, the anisotropy value was multiplied by the cosine and sine of the doubled orientation, producing weighted X and Y vector components. Weighted cosine and sine values were summed across all ROIs in the group. The mean doubled orientation (in radians) was obtained using the ATAN2 function. The doubled mean angle was halved to return to axial space and converted back to degrees. This method was validated using test inputs (e.g., 5° and 10° with equal anisotropy weights), which yielded the expected mean of 7.5°. Mean group microtubule orientations were plotted onto schematic coordinate systems to represent cell positions relative to the LRP outline using a custom Python script "Supracellular-microtubule- orientation.ipynb" (https://doi.org/10.5281/zenodo.17598920).

**Investigation of cell-level correlation between cortical SOK signal intensity and anisotropic growth**

In MorphoGraphX, numerically labelled meshes from the anisotropy measurements were used. The SOK signal image was projected onto these labelled meshes, and signal intensities were exported for each cell label. However, because the anisotropic growth calculation required an initial time point with a different set of numeric labels, the anisotropy data could only be exported using the labels from that initial time point. As a result, the numeric cell labels between the cortical signal intensity and the anisotropic growth datasets did not correspond. To address this, individual cells were manually matched between datasets based on their spatial positions. This produced a combined dataset containing both anisotropic growth and cortical signal intensity values per cell, which was then analyzed in Excel using Pearson and Spearman correlation tests.

**CarboTag fluorescence lifetime analysis**

Mean intensity–weighted fluorescence lifetimes were measured using the Leica LAS X FLIM software. Cell–cell interfaces within vascular and ground tissue precursor cells were manually outlined using the draw tool. For each tissue domain, mean intensity–weighted fluorescence lifetimes were obtained by fitting the fluorescence decay curve with a two-component exponential model. The resulting data were visualised in RStudio, and statistical significance was assessed using separate Wilcoxon signed-rank tests (script boxPlot-CarboTag.R, https://doi.org/10.5281/zenodo.17598920) .

**Turgor-induced strain simulations and extraction of SOK intensity along cell edges**

To model strain patterns in lateral root primordia, we used the vertex-based mechanical framework of Ramos et al. (2024), in which cell walls are represented as elastic one-dimensional elements arranged in a 2D lattice. The total mechanical energy of the tissue is minimised under three contributions:

1. **Elastic (stretching/compression) energy**, which depends on the Young's modulus (*E*), determines the stiffness of each wall segment against longitudinal deformation.
2. **Bending energy**, governed by the bending modulus ($k_{\mathrm{bend}}$), penalises deviations of cell corner angles from their rest configuration and captures the resistance of junctions to curvature.
3. **Pressure energy**, defined by the turgor pressure (*T*), which acts uniformly on each cell and drives expansion of the pressurised lattice.

In detail, the complete mathematical description of the system energy is given by

$$\mathcal{H} = \sum_{\mathrm{walls}} \frac{l_{0\mathrm{w}}}{2} E \left( \frac{l_{\mathrm{w}} - l_{0\mathrm{w}}}{l_{0\mathrm{w}}} \right)^2 - \sum_{\mathrm{cells}} T A_{\mathrm{c}} + \sum_{\mathrm{cells}} \sum_{\mathrm{junctions}} k_{\mathrm{bend}} \frac{1 - \cos\left(\theta_{\mathrm{c,j}} - \theta_{0\mathrm{c,j}}\right)}{\Lambda_{\mathrm{c,j}}},$$

where the subscripts $w$, $c$, and $j$ refer to a specific wall, cell and junction respectively. Wall length, $l_w$, cell area, $A_c$, junction angle $\theta_{c,j}$ and junction arclength, $\mathbf{\Lambda}_{c,j}$ are functions of vertex positions. Walls also have a rest length $l_{0_w}$ and a junction rest angle $\theta_{0_{c,j}}$, which are set to the values extracted from segmentation. For each junction, there is an angle for each cell, hence the double index, and the junction arclength is simply half the combined length of the two walls adjacent to $j$ of cell $c$. The width of the wall is taken implicitly into account within the parameters $E$ and $k_{bend}$. The energy function is then minimised w.r.t. the vertex positions using a limited-memory Broyden–Fletcher–Goldfarb–Shanno (L-BFGS) quasi-Newton method (Nocedal & Wright, 2006). In this model, each wall segment undergoes elastic deformation according to Hooke's law, while junctions resist bending due to the energetic cost of changing internal cell angles. Turgor pressure provides the driving force for wall expansion, and mechanical equilibrium is reached when the sum of stretching, bending, and pressure energies is minimised. By adjusting the three key parameters (Young's modulus, bending modulus, and turgor pressure), we simulated how tissue geometry and mechanical constraints shape the strain distribution within the primordium. The simulations were performed on meshes manually segmented from experimental images using the plasma membrane marker (pUBQ10::LTI6B:mCherry), allowing us to compare the predicted strain with the measured SOSEKI signal intensity along individual cell interfaces. To quantify SOK fluorescence intensity along cell edges, the experimental SOK images were overlaid onto the corresponding segmented mesh. For each line segment, signal intensity was estimated by sampling the fluorescence along the edge using a Gaussian convolution ($\sigma$ = 4 pixels). The resulting values were rescaled between 0 (background across all images) and 1 (maximum signal minus background across all images).

**Domain-specific strain analyses (Fig. 2).** For simulations of *pRPS5A::SOK2:YFP* primordia (stages IV–VI), the mechanical parameters were set to:

- E: 38 MPa
- $k_{bend}$: 2645 MPa
- T: 0.6 MPa

To emulate mechanical resistance from surrounding tissues, vertices located at the periphery of the mesh were held fixed (Fig. S7), whereas internal vertices remained unconstrained. The lattice was manually partitioned into three domains: provascular, boundary, and ground-tissue precursors.

**Ablation simulations and wall-type analysis (Fig. 3).** For simulations of primordia with and without ablation (stages III-V) and the correlation with *pRPS5A::SOK5:YFP signal*, cell edges were automatically classified as anticlinal or periclinal based on orientation relative to the primordia surface. Accuracy was verified by eye. Mechanical parameters were adjusted to produce strain magnitudes comparable to the experimental conditions:

- E: 300 MPa
- $k_{bend}$: 10 GPa
- T: 0.2 MPa

These higher stiffness values and reduced turgor reflect the modelling of a softer mechanical restraint.

To model ablation, we defined two sets of external vertices: a central set free to move, and a peripheral (frustrated) set that constrains the expansion of the primordium in that region. When simulating the strain distribution in the ablated case, the fraction of free-to-move vertices was increased compared to the control condition to account for the absence of the overlying tissue. The frustrated vertices have an additional energy term ½ k($\mathbf{x}$-$\mathbf{x}_0$)$^2$, with spring constant k=0.4 MPa $\mu$m, $\mathbf{x}_0$ the initial position and $\mathbf{x}$ the current one.

For all simulations, predicted strain values for each cell edge were plotted against the corresponding experimentally measured cortical SOK fluorescence intensity to assess the relationship between local mechanical state and SOSEKI polarisation.

**Quantification of SOK intensity under compressive stress (pRPS5A::SOK2/5:YFP)**

LRP length, SOK signal intensities, and plasma membrane marker signal intensities were measured in Fiji using manually drawn ROIs (for signal intensities) or lines (for LR length) in each time point. The SOK signal intensities were normalised by dividing each SOK signal intensity by the plasma membrane signal intensity and scaling it to 0-1 using linear regression. LR length was similarly scaled between values 0 and 1. The data was plotted over time in RStudio.

**Quantification of SOK intensity under compressive stress (PHS1ΔP)**

Seedlings homozygous for pRPS5A::SOK5:YFP, pCASP1::MAP4:mVenus, and pCASP1>>PHS1ΔP were transferred to mock (½MS) or DEX (½MS + 15 μM dexamethasone) medium at 6 DAG. Following 24 h gravistimulation, stage II primordia were imaged. In Fiji, cortical SOK5 and MAP4 signal intensities were measured at central cell–cell interfaces (line width 5 px), excluding the endodermis. Mean signal intensities were calculated and compared between treatments in RStudio using custom scripts (boxPlot-CarboTag.R, https://doi.org/10.5281/zenodo.17598920).
